# Global proteomic analysis of sea urchin embryos reveals dynamic changes throughout embryogenesis

**DOI:** 10.64898/2026.09.18.752731

**Authors:** Michael D. Testa, Yanbao Yu, Jia L. Song

## Abstract

The sea urchin is a long-studied model used to study fertilization and various developmental processes. While its genome and transcriptomics data are well documented, the global quantitative proteomic dynamics that complement mRNA expression during development are not available. Here we utilize an “in-cell proteomics” strategy to profile the proteomic changes in 5 developmental time points (egg, 32-cell, blastula, gastrula and larva) of the purple sea urchin, *Strongylocentrotus purpuratus*. This strategy bypasses cell lysis needed in conventional proteomic experiments, and detects over 6,000 protein groups using only 50 eggs/embryos. Over 3,000 proteins have significant changes throughout development that cluster into 6 unique expression patterns. Analysis of signaling proteins and some transcription factors involved in cell specification and differentiation align well with published literature, supporting that proteomes obtained reflect biology of the organism. Furthermore, a majority of mRNA and their corresponding protein levels are discordantly correlated across development. Overall, this study provides a comprehensive analysis of proteomic changes occurring throughout sea urchin development and establishes a powerful and sensitive high-throughput approach suitable for detecting proteomic changes with a small number of embryos.

**Summary statement:** This study establishes a novel high-throughput proteomics approach used to provide a comprehensive analysis of proteomic changes occurring throughout sea urchin development.

## Introduction

The purple sea urchin, *Strongylocentrotus purpuratus,* has been used as a developmental model organism for over a century [1]. Sea urchins are echinoderms that are closely related to chordates, containing the same major gene families as humans [2]. The phylogenetic position of the sea urchin as a basal deuterostome makes it an invaluable outgroup for assessment of evolutionarily conserved biological processes, especially in revealing the gene regulatory network (GRN) architecture and functionality using comparative genomics and transcriptomics [3–5]. Sea urchins have a high fecundity and undergo external fertilization, resulting in transparent embryos that develop over a predictable life cycle. Additionally, this organism is exceptional in withstanding experimental perturbations, such as microinjections and transplantations, making them an attractive model for studying development [4].

Previous research using the sea urchin has revealed key insights into cellular and developmental processes, such as fertilization [6], the discovery of cyclins [7], morphogenesis, and the establishment of systems-level GRNs as a means to control cell specification and differentiation [5, 8–14]. In addition, a plethora of transcriptomics of *S. purpuratus* embryos (both whole-embryo, adult tissues, and single-cell RNA sequencing) are available that allows for mapping of cell states through gene expression data at different early embryonic timepoints and tissues [15–20]. To date, physiological proteomic analyses using the sea urchin have only focused on egg activation [21], adult tissues or single tissue types, such as coelomic fluid [22], tooth matrix [23] and larval skeletal matrix [24]. Other studies utilized proteomics to analyze changes under infection stress in adult coelomocytes [25], temperature stress in adult tube feet [26], and ultraviolet radiation stress in embryos [27]. However, quantitative proteomic changes that complement transcript data throughout early sea urchin embryogenesis are absent. In addition, the gold standard gene perturbation approach using the sea urchin embryos involves technically challenging microinjections that result in relatively few embryos. Thus, sensitive detection of proteomic changes in experimentally perturbed embryos has not been feasible or practical. Consequently, we have a strong need for quantitative proteomics approach that is sufficiently sensitive to detect proteomic changes in a relatively small number of experimentally manipulated embryos.

Bottom-up proteomics typically processes proteins upon cell lysis and protein extraction with chemical or mechanical disruptions, followed by precipitation or cleanup before achieving trypsin-compatible digestion conditions. However, the above preparations are associated with lengthy and complex procedures as well as potential sample losses, which are detrimental to analysis of quantity-limited samples. Recently, we developed a simple yet high-sensitivity “in-cell proteomics” strategy, where the proteins are digested directly inside of methanol-fixed cells [28]. The on-filter in-cell (OFIC) digestion approach bypasses traditional multi-step sample pretreatment steps, prevents material loss while maintaining high digestion efficiency and quantitative reproducibility, offering a promising alternative to low-cell/low-input proteomics. The robustness and wide applicability of the method has been successfully demonstrated in mammalian, bacterial and fungus cell line samples [29–31], plant tissues [32], as well as embryonic samples from nematodes and frogs [33, 34].

Here, we present for the first time using this highly sensitive ‘in-cell proteomics’ approach to provide a valuable resource documenting a comprehensive analysis of proteomic changes from eggs to larval stages and reveal major proteomic shifts and patterning during early embryogenesis.

## Methods

### Animals, Embryo Culture and Fixation

Adult *Strongylocentrotus purpuratus* (Sp) were collected from the California coast (Fisherman Marketing Association, Lakeside, CA or Marinus Scientific, LLC, Long Beach, CA). All animals and embryonic cultures were incubated at 14°C. Newly fertilized zygotes from three adults were cultured in filtered seawater (FSW) obtained from the Indian River Inlet (University of Delaware) in a 24-well plate at 14°C. 50 eggs and embryos at different developmental time points (Fig.1). Samples were collected, spun down, and fixed with 450µL of cold 100% methanol and stored at - 80°C until proteomic analysis.

**Figure 1.**
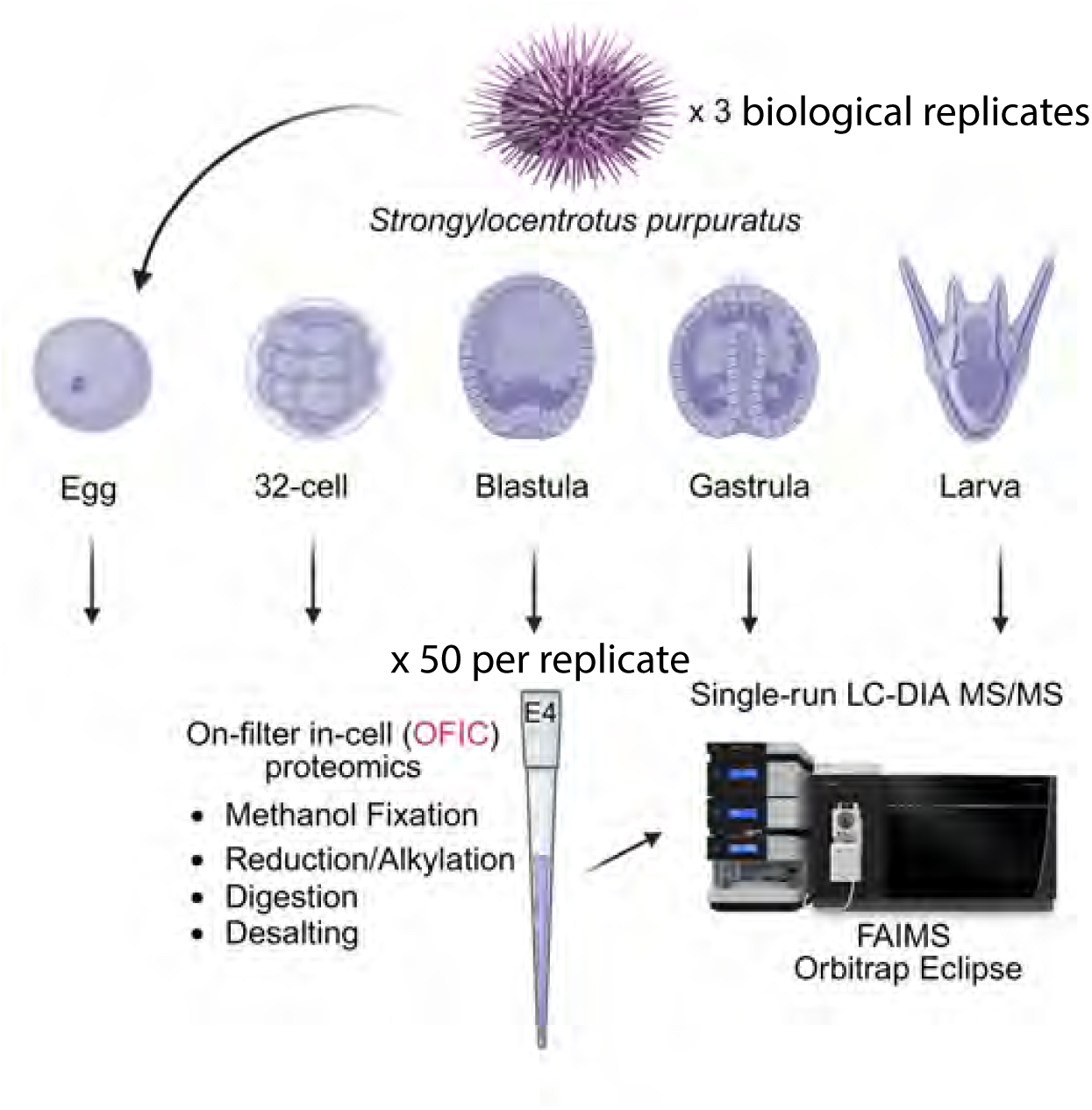
Workflow of the in-cell proteomics approach for sea urchin eggs/embryo analysis. Embryonic samples derived from egg, 32-cell, mesenchyme blastula, gastrula, and larval stages were collected and subjected to methanol fixation, reduction and alkylation, protein digestion and peptide desalting in E4tips. 50 eggs/embryos from each stage were used for each digestion experiment (3 bioreps). Mass spectrometric analysis was performed using Orbitrap Eclipse with FAIMS Pro Interface in DIA mode.

### Embryo sample preparation for proteomics

Sea urchin embryos from different developmental stages were collected, transferred to low-binding microtubes, and incubated with pure methanol on ice for 10 min. The embryos were then subjected to OFIC digestion following a protocol described recently [33]. We utilized 50 embryos per digestion and performed three digestion experiments per developmental stage. Briefly, the methanol-fixed embryos were transferred to E4tip XL (CDS Analytical, Oxford, PA), spun at 1,500 *x* g for 2-3 min to discard the flow through. The samples were then subjected to reduction and alkylation by adding final concentration of 10 mM Tris (2-carboxyethyl) phosphine (TCEP) and 40 mM chloroacetamide (CAA) in 200 μL of 50 mM triethylammonium bicarbonate (TEAB), and incubated at 45⁰C for 10 min. The tips were spun, washed once with 200 μL of 50 mM TEAB, and then transferred to new collection tubes.

For protein digestion, 2.0 μg of trypsin and 2 mL of 50 mM TEAB buffer were added to the E4tips, which were then incubated at 37⁰C for 16-18 hours with gentle shaking (300 rpm). After digestion, 5 μL of pure formic acid was added to the E4tips, and incubated under RT for 3-5 min. The tips were then centrifuged at 1,500 *x* g for 10 min, followed by a wash step with 200 μL of 0.5% acetic acid in water. The E4tips were transferred to new collection tubes and eluted sequentially with 200 μL of 0.5% acetic acid in 60% ACN and another 200 μL of 0.5% acetic acid in 80% ACN. The elution was dried in SpeedVac and stored under -80⁰C until further analysis.

### LC-MS/MS analysis

The LC-MS/MS acquisition was performed using an UltiMate 3000 RSLCnano system in combination with an Orbitrap Eclipse mass spectrometer and FAIMS Pro Interface (Thermo Scientific). The trap column was PepMap100 C18 with dimensions of 300 μm × 2 mm, and a particle size of 5 μm (Thermo Scientific). The analytical column was PepMap100 C18 (50 cm × 75 μm i.d., 3 μm particle size; Thermo Scientific) flowing at 250 nL/min. A linear LC gradient was applied from 1% to 25% mobile phase B (0.1% formic acid in acetonitrile) over 125 min, followed by an increase to 32% mobile phase B over 10 min. The column was washed with 80% mobile phase B for 5 min, followed by equilibration with mobile phase A (0.1% formic acid in water) for 15 min. For MS analysis, the data-independent acquisition (DIA) mode was used. In brief, the detector type was Orbitrap with a resolution of 120,000 for full scan; Precursor MS range (m/z) was 380–980; AGC target was Standard; Maximum injection time mode was Auto. For DIA MS/MS analysis, the Isolation mode was Quadrupole; DIA Window type was Auto and Isolation Window (m/z) was 8 with an overlap of 1 m/z; activation type was HCD with fixed collision energy mode (30%); the Detector Type was Orbitrap with a resolution of 30,000; Normalized AGC target (%) was 800, and the Maximum injection time mode was Auto; the Loop Control was 2 s. For FAIMS compensation voltages (CV) setting, a 3-CV combination (−40, −55, and −75) was applied.

### Bioinformatics and data analysis

The MS raw data were processed using Spectronaut software (version 19.5) and a library-free DIA analysis workflow with directDIA+ and the *S. purpuratus* proteome obtained from UniProt Knowledgebase (taxonomy ID: 7668, 35,496 sequences). Detailed settings in Spectronaut included: Trypsin/P as the specific enzyme; peptide length from 7 to 52 amino acids; allowing 2 missed cleavages; toggle N-terminal M turned on; Carbamidomethyl on C as fixed modification; Oxidation on M and Acetyl at protein N-terminus as variable modifications; False discovery rates (FDRs) at PSM, peptide and protein level all set to 0.01; Quantity MS level set to MS2, and cross-run normalization turned on. Bioinformatics analyses including t-test, correlation, and clustering analyses were performed using Perseus software (version 1.6.2.3), and GraphPad Prism (version 11.0) unless otherwise indicated. Cellular component analysis is performed with DAVID Bioinformatics [35, 36]. Fig.4A was generated with the ClusterGVis package in R (Zhang et al., 2026) (https://github.com/junjunlab/ClusterGVis). Figs.7C,S1C density scatter plots were generated with ggplot2 and ggpointdensity in R (Wickham, 2016).

### Relative Protein and mRNA Expression Graphs

All mRNA transcripts (TPM+1) were taken from Echinobase, Log_2_ transformed and normalized to the egg. Transcripts from 0hpf (egg), 10hpf, 24hpf (Blastula), 48hpf (Gastrula) and 72hpf (Larva) were used. Note that 10hpf transcriptomic data was used to match our 32-cell stage proteomics data at 6hpf. In cases where a protein with a corresponding gene has more than one transcript listed in the Echinobase (genomic database for *S. purpuratus*), we presented all versions. Relative protein levels (Z-score) were averaged across 3 replicates and normalized to the egg. Error bars represent Standard Error of the Mean (SEM) in all protein expression graphs (Figs.5-7).

## Acknowledgements

We thank Veronica Hinman and Nicholas Christodoulides (Whitney Laboratory for Marine Bioscience, University of Florida) for providing matched UniProt accession numbers to Entrez gene IDs *en masse*.

## Results and Discussion

### Quantitative assessment of the in-cell proteomics of sea urchin embryos

To investigate the dynamic changes of early sea urchin embryogenesis, we adopted a tip-based single-vessel “in-cell proteomics” strategy, which eliminates cell lysis and protein extraction and integrates all of the treatment steps into the same microchamber, thus avoiding the multi-step transfers and minimizing surface adsorption and sample loss [28]. In this study, we pooled 50 sea urchin eggs or embryos for each in-cell digestion experiment. Overall, this dataset reported the identification of 7,261 non-redundant protein groups upon mapping the MS data to the *S. purpuratus* UniProt Knowledgebase (35,496 sequences), averaging approximately 6,600 proteins across the five developmental states (Fig.2A; Table S1). We further examined our MS data against the Echinobase (38,439 sequences), another established and centralized knowledgebase for the echinoderm community [18]. We noticed a largely 10-20% increase of protein identifications, and 10-30% increase of peptide hits (Fig.S1A), leading to a total of 8,805 proteins (Table S1). Both results represent the largest *S. purpuratus* proteome with experimental evidence reported to date. For downstream enrichment and pathway analysis, we focus on the UniProt dataset, as its identifiers are readily compatible with some of the commonly used bioinformatics tools.

**Figure 2.**
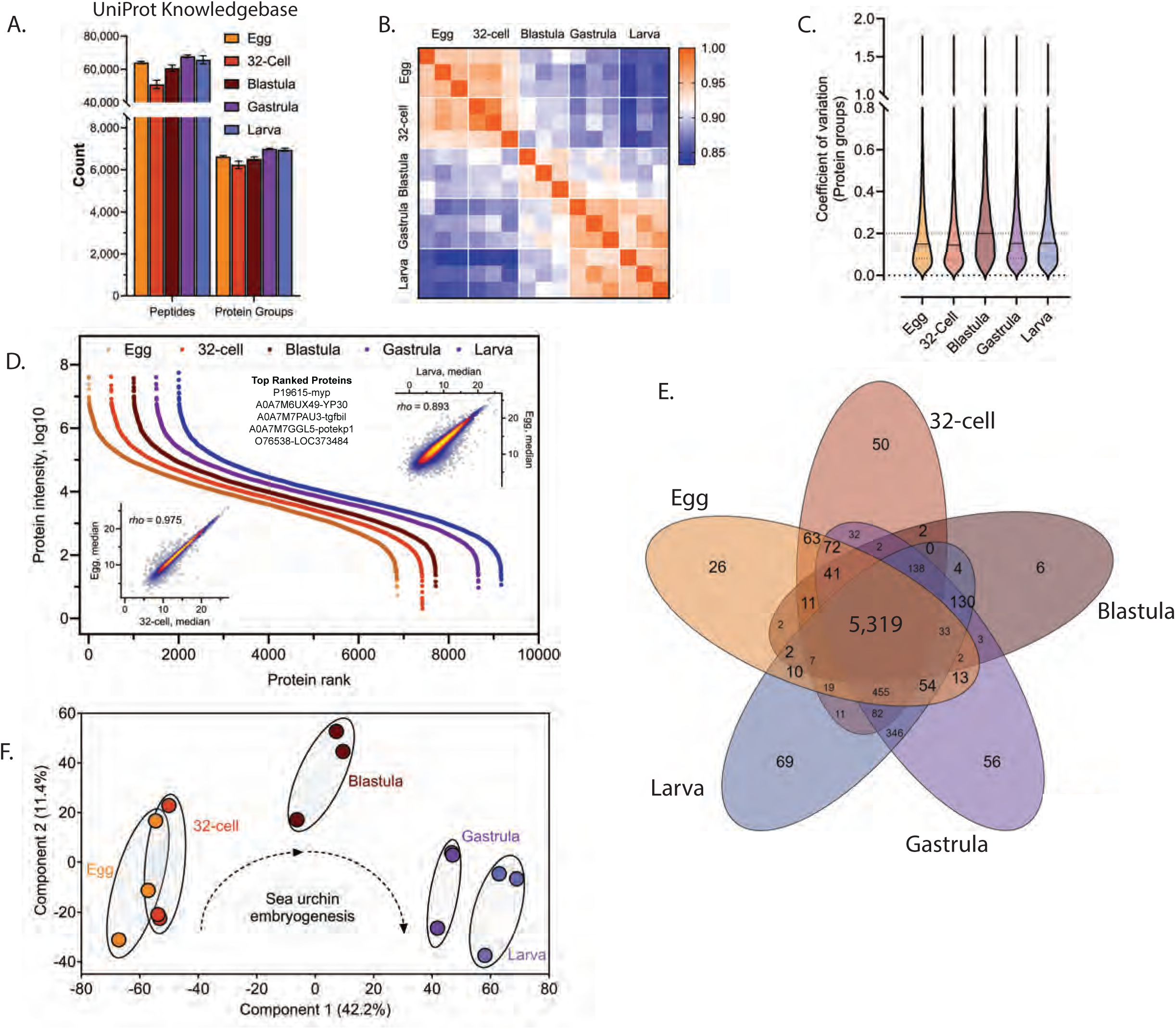
Evaluation of the in-cell proteomics approach. (A) The total identification of peptides and protein groups across 5 developmental stages (egg-larvae) is shown. Error bars represent standard deviation among 3 bioreps. (B) Pearson’s correlation analysis across 5 developmental stages is shown. (C) Coefficient of Variation (CV) of protein groups at each developmental stage is shown. Horizontal solid lines indicate the median value (D) Protein rank plot across five developmental stages is depicted. Top 5 proteins at each stage are indicated in the graph. The inner panels show two representative correlation plots between egg and 32-cell, and between egg and larva, respectively. The Spearman correlation values are depicted on the top left of the panels. (E) Venn diagram of shared or unique proteins is shown across five developmental stages. (F) Schematic of principal component analysis (PCA) of 3 bioreps in each developmental stage is shown.

We validated the quality of the proteome profiles of 5 developmental timepoints (egg-larva) Regarding variability, the in-cell approach achieved excellent correlation (Pearson ∼ 0.95) between the biological replicates, and the median coefficient of variation (CV) of the protein groups were 20% or lower for each stage (Fig.2B-C). These data are consistent with the performance of the in-cell digestion approach in several other organisms and human cell lines [29–34], suggesting its robustness for global quantitative proteomics analyses. We further analyzed proteome-wide similarities and differences among the five stages. Results indicate that the sea urchin proteome spans nearly eight orders of magnitude at every one of the five stages (Fig.2D and Fig.S1B). Such a wide and dynamic range highlights the enormous differences of the sea urchin cellular protein abundances and underscores the analytical challenges of detecting rare or low-abundance proteins in this model organism. As expected, major yolk protein (P19615-myp) is among the top ranked hit from our analysis along with other yolk proteins A0A7M6UX49-YP30 and A0A7M7PKP5-Apob (vitellogenin). Myp is a 180-kDa glycoprotein that accounts for 10-15% of the entire cellular proteome [37] and acts as the primary nutrient source for the developing embryo in both vertebrate and invertebrate species. In-cell proteomic analysis in *Xenopus* [38] and 2D-LC-tandem MS (MS/MS) and LC-CZE-MS/MS in *Danio* [39] revealed similar results with highly ranked yolk proteins. In all cases, eggs and embryos were not stripped of yolk, which could lead to masking non-yolk proteins. A Venn diagram demonstrates that the majority of proteins (89.1% or 6,433/7,261) are shared by all the developmental stages (Fig.2E and Table S1), indicating substantial proteomic similarities during early embryogenesis [40]. A principal component analysis (PCA) illustrated largely distinct developmental proteomes, although marginal overlaps were seen between the stages of egg and 32-cell (cleavage) stage (Fig.2F). This likely reflects the residual maternal proteins of the egg that are still present in the cleavage stage embryos at 6 hours-post-fertilization (hpf). Of note is the clear spatial distinction the blastula-stage proteins exhibit when compared to all other stages collected in the PCA plot (Fig.2F). Consistent with our data, transcriptomic analysis of the green sea urchin (*Lytechinus variegatus*; Lv) reveals that the hatched blastula stage accounts for the largest variation in the Lv transcriptome and is distinct from all other developmental time-points analyzed [20]. This indicates that embryos exhibit a drastic proteomic transition from cleavage stage into blastula and thereafter into gastrulation and larval development.

### Greatest proteomic change occurs in the blastula stage

A cellular component analysis using DAVID Bioinformatics [35, 36] of total 7,261 identified proteins (Fig.3A and Table S1) indicates that the majority of proteins detected reside in the cytoplasm (cytosol plus organelles) (20.49%), with the next top components being the nucleus (17.94%) and cytosol (7.21%). A broad distribution of cellular components was detected that includes the mitochondria (5.39%), endoplasmic reticulum / membrane (5.47%), Golgi (2.23%) and cytoskeleton (3.04%). The extensive coverage of cellular components observed in the sea urchin is consistent with that of the frog and nematode during early embryogenesis using the E4-technology [38, 41] and the zebrafish using 2D-LC-tandem MS (MS/MS) and LC-CZE-MS/MS [39].

**Figure 3.**
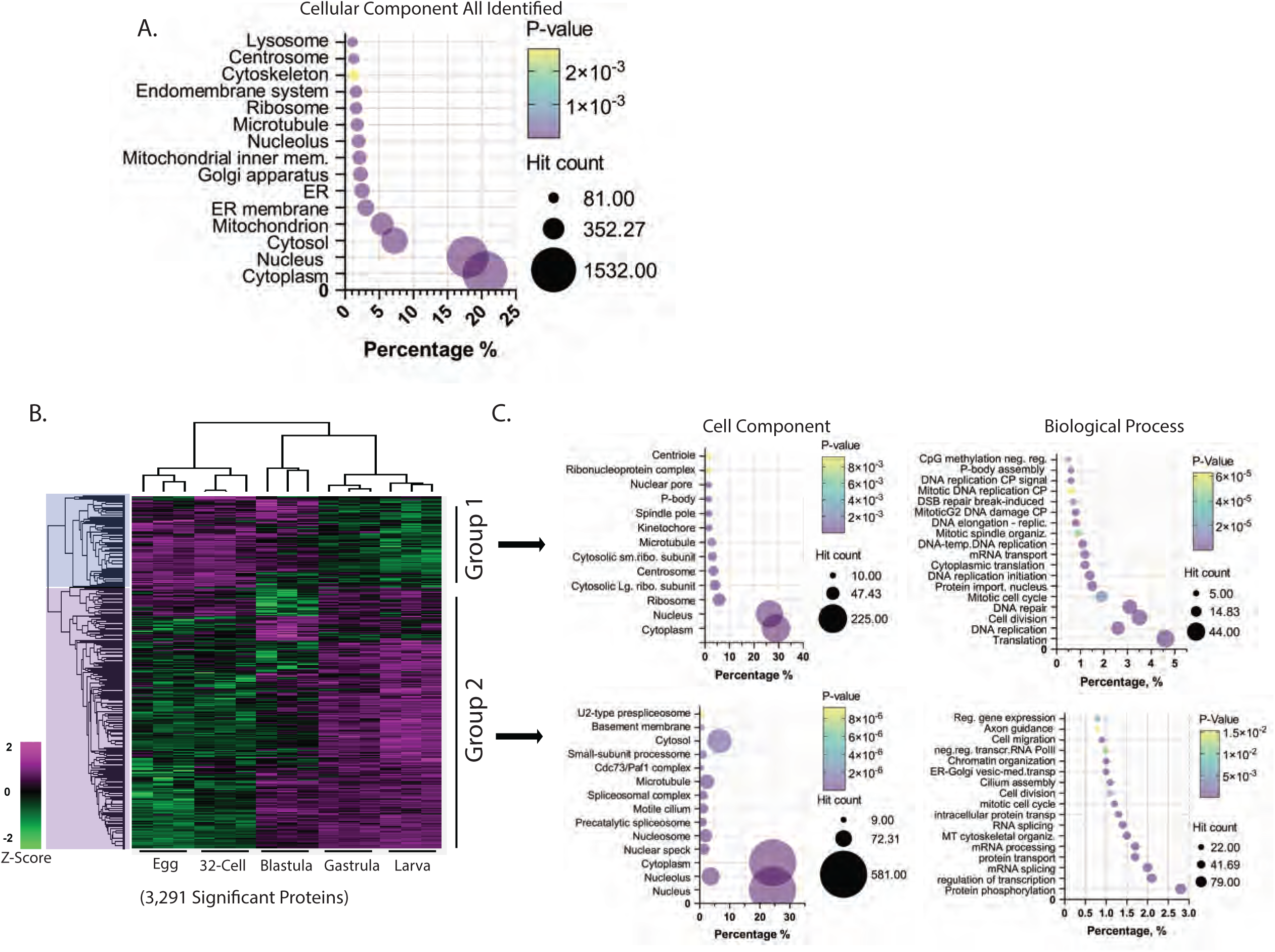
Overview of significantly changing proteome across development. (A) DAVID bioinformatics [36] analysis of all identified proteins is used to classify proteins (7,261 protein groups) across development into cell component and biological processes. (B) Multi-sample test (ANOVA, permutation False Discovery Rate (FDR)=0.05) among the five developmental stages reveals 3,291 proteins to be significantly different in protein levels throughout development. Heat map indicates relative protein expression changes (Z-score) from egg-larva. Groups 1 and 2 of 3,291 significant proteins are highlighted. (C) DAVID bioinformatics analysis of 3,291 significantly changed proteins across development is performed for cell component and biological process.

Multi-sample test (ANOVA, permutation False Discovery Rate (FDR) = 0.05) among the five developmental stages indicate that, of the total 7,261 protein groups, 3,291 proteins (45%) displayed significant difference in their levels among the five developmental stages, indicating that a majority of proteins (55%) remain statistically unchanged throughout development (Fig.3B and Table S1). The ⁓55% of proteins that remain unchanged may be due to their housekeeping functions necessary through all stages of development, or that the threshold in this approach is limited in discerning subtle changes in the proteome. The 45% of significantly different proteins (3,291 proteins) that were detected are grouped into two major distinct groups based on their general protein expression trends (Fig.3B). This includes the egg-32-cell (Group 1 with 847 proteins) and gastrula-larva stages (Group 2 with 2,444 proteins). One apparent observation is that the blastula proteome has a high variability in protein levels (Fig.3B), indicating a potential significant overhaul of maternally derived proteins during blastulation. Group 1 is enriched with highly maternal proteins involved in translation, DNA replication/repair and mitosis (Fig.3C) which reflects the rapid and coordinated cell divisions immediately following fertilization that primarily rely on translation of maternally stored mRNAs [42–45]. This is further supported with cell component analysis of Group 1, indicating the ribosome and ribonucleoprotein complex protein enrichment (14.5%) (Fig.3D,E). Group 2 is enriched with proteins relating to RNA synthesis/metabolic processes, such as chromatin organization, regulation of transcription and mRNA splicing (Fig.3C) and are highly increased post-blastulation. Our data likely reflect the transition the embryo incurs while activating its genome post early-cleavage into the blastula stage, coinciding with the larger, more pronounced wave of zygotic transcription that occurs during the blastula stage [45–47]. Cell component analysis corroborates this observation, as many proteins shift from ribosome residency in the early cleavage stage to residing mainly in the nucleus (36.96% compared to 28.9%) (Fig.3D). Taken together, these results indicate that the OFIC approach has the capability of detecting a broad range of cellular components during early embryogenesis and reveal that the blastula stage as a developmental stage of significant proteomic landscape transitioning, similar to prior transcriptomic data [20].

### Hierarchical clustering of proteins based on their expression patterns

From our proteomics analysis, the 3,291 significantly different proteins across the five developmental stages are further clustered into detailed expression patterns (Z-score enrichment), using the ClusterGVis package in R (Fig.4) [48]. The blastula stage is a period in development when the embryo begins major waves of zygotic gene activation and cells start to differentiate and the embryo undergoes major morphogenetic changes. We identify 6 clusters of expression patterns: proteins in Cluster 1 (946 proteins) and cluster 4 (1,002 proteins) remain relatively low until the gastrula and larval stages; however, proteins in Cluster 4 begin to accumulate into the blastula stage. Proteins in Cluster 3 (468 proteins) and Cluster 5 (368 proteins) are high in the eggs and low in the gastrula and larval stages. However, proteins in Cluster 3 remain enriched in the blastula, whereas proteins in Cluster 5 do not. Proteins in Cluster 2 (207 proteins) and Cluster 6 (300 proteins) have almost inverse protein levels, where Cluster 2 proteins have high enrichment in the blastula stage only, and Cluster 6 proteins have very low levels during this stage.

**Figure 4.**
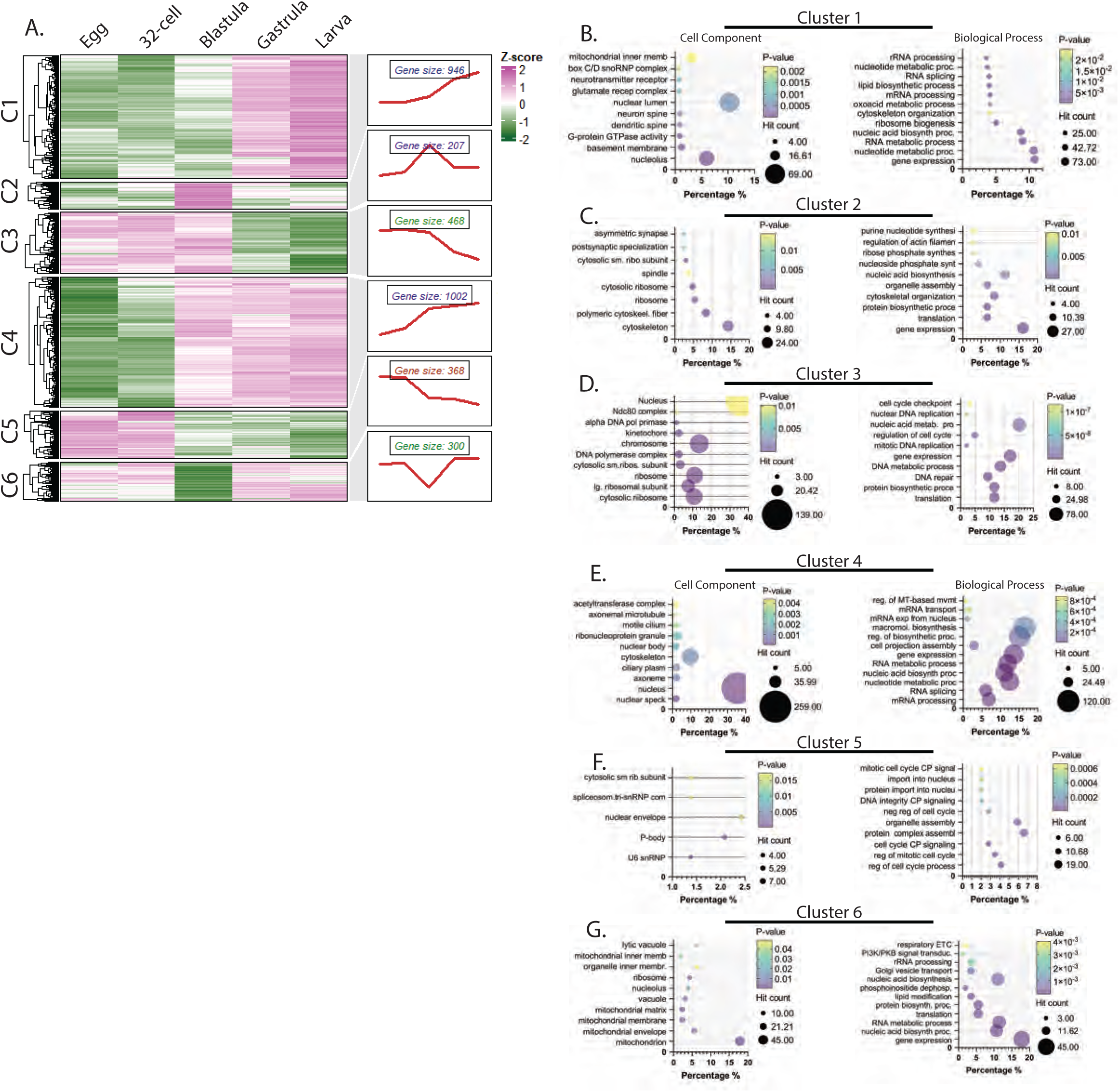
Hierarchical clustering reveals specific protein trends. (A) Proteins (3,291) that have levels significantly different throughout development were subjected to ClusterGVis R package to analyze expression patterns. 6 clusters are represented (B-G) DAVID Bioinformatics analysis of Clusters 1-6 are shown for cell component and biological process.

Further analysis of Cluster 2 proteins indicates an enrichment with proteins involved in gene expression (16.07%), nucleic acid biosynthesis (11.31%), RNA metabolism (8.33%), and cytoskeletal organization (8.33%) (Fig.4B). Cluster 4 proteins are primarily related to regulation of RNA metabolism (11.92%), metabolic processes (16.44%), biosynthesis (15.07%), and nucleic acid metabolism (13.15%) (Fig.3C). Clusters 2 and 4 (combined 36.74%) proteins are at low levels in the egg and 32-cell stage and steeply increased in levels in the blastula stage (Fig.4A,C,E). However, proteins in Cluster 2 are unique in that these proteins are only at high levels in the blastula stage and decreased thereafter. Some notable proteins in Cluster 2 include Disheveled, Sp-Ef1a, eif3j, etf1, exosc1, exosc2, mettl1, and a large number of ribosomal proteins (rps23, rps19, rpl37a, rpl23, rpl11, LOC578794, LOC115918936 and LOC579026). Exosc1 and Exosc2, whose main function is 5’ → 3’ RNA degradation and processing, are required for early embryonic development in mice [49].

Interestingly, they play different roles, where Exosc1 null embryos are developmentally delayed and fail to gastrulate and Exosc2 null embryos are lethal during peri-implantation stages [49]. The increased protein level of Exosc1 and Exosc2 preceding gastrulation may indicate their conserved function in development. The sudden increase of ribosomal and translation-related proteins could be needed to compensate for the increase in newly transcribed mRNAs in the blastula stage that follow the major wave of zygotic genome activation [45, 46]. Expectedly, one of the key hatching enzymes (Sp-He2 LOC752524), which is a matrix metalloprotease that promotes the release of the blastula from the fertilization envelope, has highest protein level in Cluster 2 [50, 51]. These hatching enzymes are significantly upregulated during the blastula stage and at no other time point, reflecting their functions during blastulation.

Cluster 4 proteins, which are gradually increased into blastula and thereafter, include a large number of mRNA binding and mRNA processing proteins. Interestingly, a large number of these proteins are involved in RNA splicing (LOC115921160, LOC584228, sf3b1, sf3a3, sf3a1, sart1, LOC582113, LOC763151, LOC588026, prpf4b, prpf31, cwc15, crnkl1, cactin and aar2), potentially reflecting the increase in zygotic gene activation and transcription during the blastula stage and later development (Fig.4 and Table S1). Many metazoan embryos undergo a period of maternal clearance, in which maternal proteins and RNAs are cleared from the embryo to allow embryonic cell specification and differentiation [52–54]. microRNAs have been shown to be critical to mRNA clearance in the frog [55], fly [56] and zebrafish [57], mainly by deadenylating transcripts. Cluster 4 proteins include Ago1, seawi, and piwil1, members of the RNA induced silencing complex (RISC), as well as proteins involved in exosome (LOC762762 and dis3) and deadenylation (cnot6l and LOC591400), which may be upregulated during the blastula stage to facilitate mRNA degradation and clearance (Fig.4 and Table S1).

Cluster 6 proteins are dramatically decreased in the blastula stage but have increased protein levels in the gastrula and larval stages (Fig.4A,G). Proteins in this cluster are enriched in mitochondria (17.79%), involved in gene expression (17.79%), involved in nucleic acid biosynthesis (11.07%), and protein biosynthetic processes (5.53%). The larger groups of mitochondrial genes in Cluster 6 includes mitochondrial ribosomal proteins (mrps22, mrps17, mrps12, mrps10, mrpl9, mrpl44, mrpl24, dimt1, LOC764162, LOC115919962 and LOC588867), cytochrome c oxidase subunits and assembly components (LOC115918811, LOC585298, LOC591899, cox15) and oxidoreductase subunits (LOC754255, ndufs8, ndufaf1). Little has been reported about the relative metabolic pathways used at differential phases of early development. In the sea urchin, inhibition of fatty acid synthesis and glucose metabolism result in dose-dependent gastrulation failure [58]. In addition, it has been shown that mitochondrial metabolism is essential for early embryonic development, because the embryo undergoes rapid cell divisions that require a constant supply of ATP through oxidative phosphorylation, tricarboxylic acid cycle (TCA), and electron transport chain [59–61].

### Increased levels of transcription factors are observed during and post-blastulation

Since transcription factors (TFs) play such critical roles in development, we examine this group of proteins across the five developmental stages (Fig. 5A). Using the GO term “DNA-binding transcription factor activity” (GO:0003700) and the GO term “transcription coregulator activity” (GO:0003712) [35], we are able to detect 222 unique UniProt accessions within our total proteomic dataset with GO:0003700 and 81 under GO:0003712 (Fig.5 and Table S1). TFs are experimentally difficult to detect due to their low abundance in the cell and are often masked by other abundant proteins (Nagore et al., 2013; Simicevic & Deplancke, 2017). To account for potential redundancy, we mapped these combined 303 accessions back to 243 annotated genes in the sea urchin genome [18]. The total number of UniProt accession TFs under GO:0003700 were further classified by domain (PFAM). Results indicate the largest number of proteins being C_2_H_2_ Zinc finger (Znf) domains (30.49%), followed by Homeodomain-like Superfamily (9.76%) (Fig.5B). This result is consistent with C_2_H_2_ zinc finger genes being the most abundant zinc finger protein domains in eukaryotes [62].

**Figure 5.**
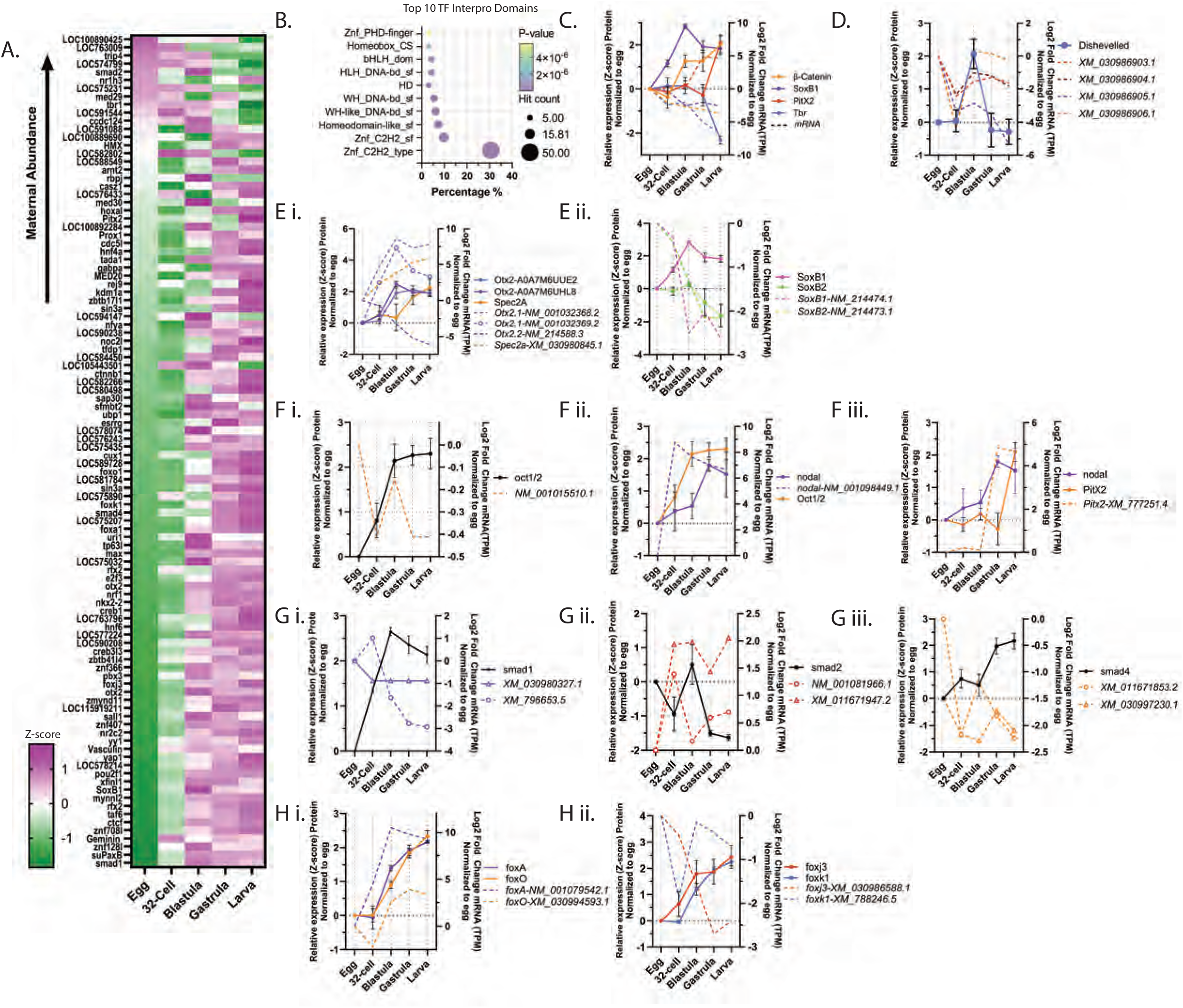
Analysis of mRNA-Protein correlations of TFs and signaling pathway components. (A) TFs and TF coregulators that are significantly different in protein levels across development are graphed in a heatmap, in order of high-low maternal protein abundance. (B) DAVID bioinformatics analysis of TF (GO:GO:0003712) characterized by InterPro domain [36] is performed. (C) *β-catenin*, *SoxB1*, *PitX2* and *Tbr* mRNAs (Log_2_TPM) and proteins (Z-score) normalized to the egg are plotted. Proteins are denoted by solid lines and their corresponding mRNAs are of the same color denoted by dashed lines. (D) Four *Dishevelled* transcripts and Dishevelled protein normalized to the egg are plotted. (E) *Otx2.1, Otx2.2* and *Spec2a* mRNAs and Otx2, Spec2a proteins normalized to the egg are plotted. (Fi) *Nodal* mRNA and Nodal, Oct1/2 proteins normalized to the egg are plotted. (Fii) *Nodal* mRNA and Nodal and Oct1/2 proteins normalized to egg are plotted. (Fiii) *Pitx* mRNA and Nodal and Pitx2 proteins normalized to the egg are plotted. (Gi-iii) *Smad1/5/8*, *Smad 2/3* and *Smad 4* mRNAs and proteins normalized to the egg are plotted. (Hi-ii) *FoxA*, *FoxO*, *FoxJ3* and *FoxK1* mRNAs and proteins normalized to egg are plotted. Proteins are denoted by solid lines and their corresponding mRNAs are of the same color denoted by dashed lines for all graphs.

Previous analysis of zinc finger genes using transcriptomics revealed 377 genes in the sea urchin genome, which is about half found in mice or human [63]. The relative number of TF hits within our proteomic dataset is comparable to *Xenopus* early embryos using E4 technology (331 TFs) [38]. In this study, we are able to identify a significantly larger number of TFs that other proteomic methods. For example, 73 TFs were detected in early zebrafish embryos, using 2D-LC-tandem MS (MS/MS) and LC-CZE-MS/MS [39].

We then asked if we could detect subtle changes in TF levels across early development. Of the 303 TFs and TF coregulator accessions identified, 103 unique TF genes mapped in Echinobase were significantly different (ANOVA, FDR=0.05) among the 5 developmental stages (Fig.5A-F and Table S1). Most significant TFs have relatively low protein levels in the egg compared to blastula/post-blastula, where the majority of TFs proteins begin to increase (Fig.5A). Increased TFs coincide with cell specification and differentiation events that occur during blastulation. A closer examination of maternal proteins reveals presence of TFs, transcription co-activators, and signaling molecules that drive early embryonic development and establish germ layer specification, and initiate zygotic genome activation in the early embryo [64]. Some of the specific key regulatory proteins are discussed below.

### Regulatory proteins of early embryonic axis and germ layer specification have significantly different levels throughout development

In the sea urchin, animal/vegetal axis specification is determined early in the egg with the localization of mRNA and protein determinants [65, 66]. Specifically, the maternal scaffolding protein Disheveled (Dvl) in the vegetal cortex of the egg stabilizes β-Catenin, allowing it to enter the nucleus of vegetal micromeres/macromeres to specify the endomesoderm. We identified Disheveled and 2 unique β-Catenin proteins with different expression patterns (A0A7M7HIL0 and A0A7M7N400). The relative protein expression of β-Catenin-A0A7M7HIL0, the more widely known co-transcriptional activator, decreases from the egg into the 32-cell stage and increases in the blastula and larval stages (Fig.5C). Interestingly, their mRNA transcript counterpart exhibits the opposite trends, in that β-Catenin-A0A7M7HIL0 protein greatly increases from 32-cell stage to blastula stage, but its mRNA steadily decreases after the 32-cell stage, reflecting that its translation is most robust between the cleavage and blastula stages (Fig.5C). Disheveled, which is a cytoplasmic Wnt pathway effector that restricts β-Catenin to the vegetal blastomeres, peaks sharply at blastula stage, and subsequently falls dramatically in later development (Fig.5D). It has been reported that the sea urchin has 4 unique isoforms of Dvl (*Sp*Dvl5a, 1, 4a and 4b) that all share >99% sequence similarity [67]. Our method was therefore unlikely to differentiate among these forms with proteomics, as 4 unique UniProt identifiers were mapped to the same gene locus (Fig.5D). However, the curated transcripts for *Dvl* exhibit increased expression in the blastula stage, with transcript *XM_030986906.1* having the greatest fold change increase that corresponds to our identified Dvl protein (Fig.5D).

The β-catenin/Wnt pathway, in turn, activates the endomesoderm GRN [68]. Otx2 (orthodenticle homeobox 2), downstream of β-Catenin, is a main activator of other early endodermally-expressed TFs [69, 70]. Our proteomics analysis indicates that two Otx2 (Orthodenticle homeobox 2) proteins (A0A7M6UHL8 and A0A7M6UUE2) have significantly different levels of protein across development (Fig.5Ei). These proteins are different in length and correspond to the different isoforms of Otx2 in the sea urchin from 3 distinct transcripts synthesized via alternative RNA splicing [71]. *Otx2.1 NM_001032368.2* and *NM_001032369.2* produce a 295aa protein (A0A7M6UUE2), while *Otx2.2 NM_214588.3* corresponds to the 371aa protein A0A7M6UHL8 (Fig. 5Ei). Both of the identified Otx2 proteins were shown to increase ⁓2-fold from cleavage stage into blastula (Fig.5Ei), while remaining steadily expressed into the larval stage. Otx2 is necessary for the activation of *Endo16* in the vegetal plate and for the activation of *Spec2a*, a calcium-binding protein belonging to the troponin C superfamily, in the aboral ectoderm [72]. As expected, Otx2 protein level peaks before the gradual increase of Spec2a protein in our dataset, reflecting Otx2 in activating Spec2a in the aboral ectoderm [72] (Fig.5Ei).

The animal-half inducing TFs, such as SoxB1, are restricted in the animal pole as development progresses to activate ectodermal GRNs [73]. SoxB1 is ubiquitously expressed and restricted to the animal blastomeres when nuclear β-catenin becomes restricted in the vegetal pole of the embryo at the 16-cell stage [73–75]. SoxB1 and its paralog SoxB2 are critical in the establishment of oral/aboral ectoderm specification and neural development [74]. We were able to detect both SoxB1 and SoxB2 in our proteomic analysis; however, SoxB1 was the only protein detected as having significantly different levels of protein among the 5 developmental stages (Fig.5C and Table S1). The relative protein and mRNA levels of both paralogs are depicted (Fig.5Eii). SoxB1 protein expression peaks at the blastula stage and levels off throughout gastrula and larva stages, likely reflecting its role in gastrulation [75] and neural specification in the early ectoderm and late endoderm of the gastrula and larval stages [74]. Interestingly, SoxB1 mRNA decreases, coinciding with accumulation of its protein throughout development, while SoxB2 protein and mRNA decrease post-blastula stage (Figs.5Eii). Thus, levels of animal/vegetal axis specification proteins such as *β*-catenin, Dvl, Otx2, Spec2a, SoxB1 detected by this proteomics approach accurately reflect known roles that these proteins play during early development.

The key TGF-β superfamily signaling molecule, Nodal, in early development is involved in the secondary, dorsal/ventral axis formation of the embryo, as well as specification and differentiation of immune cells [76–83]. Nodal is positively regulated by the maternal factor Oct1/2 [84]. *Nodal* mRNA is highly expressed up until the 32-60 cell embryo in the prospective ventral ectoderm, where it activates Pitx2 and Gsc (Goosecoid) and represses the expression of aboral specific genes, such as coquillette/Tbx2/3, resulting in specification of the oral ectoderm [77, 85, 86]. We were able to identify Oct1/2 and Pitx2 among the significantly different proteins among the 5 developmental stages (Fig. 5Fi-iii). While Nodal was not among proteins that are significantly different in expression throughout development, its protein was identified. We observed differences in *Nodal* mRNA and protein level (Fig.5Fii), in which the *Nodal* mRNA expression peaks in the cleavage stage (32-cell), whereas the Nodal protein level peaks in the gastrula stage (Fig.5Fii). As expected, protein level of Oct1/2, which positively regulates *Nodal* expression, rises rapidly from 32-cell to blastula, simultaneously with the rise in *Nodal* mRNA and preceding Nodal’s peak protein level in the gastrula stage (Fig.5Fii). *Pitx2* mRNA expression, which is activated by Nodal [87], increases during gastrulation, concurrently with peak of Nodal protein expression (Fig.5Fiii). *Pitx2* mRNA is expressed at the gastrula stage in the ectoderm near the tip of the archenteron in the right coelomic pouch [81, 88]. Our data suggest that the relative Pitx2 protein expression peaks in the larval stage (Fig.5Fiii), where it has been shown to function in the left/right asymmetry and contributes to adult rudiment formation [81, 88, 89]. Thus, the levels of Oct1/2, Nodal, and Pitx proteins are consistent with the known role of Nodal in the secondary, dorsal/ventral axis formation of the embryo.

Additional key components of the TGF-β superfamily include downstream prototypical Smad proteins, involved in early sea urchin development [90]. We identified Smad 1 (Smad1/5/8), Smad 2 (Smad2/3) and Smad 4 in our analysis as being significantly different among the 5 stages of development (Fig.5G and Table S1). Smad1 is activated by BMP signaling and has been shown to regulate left-right asymmetry in the developing embryo [81, 91–93]. Smad1 is primarily phosphorylated on the dorsal side of the embryo and partly in the dorsal primary mesenchyme cells (PMCs) at the beginning of late blastulation, contributing to dorsal skeletal patterning downstream of BMP signaling [93–95]. Our analysis indicates that relative Smad1 protein level peaks at the blastula stage (Fig.5Gi) and levels off into the larval stage. Phosphorylated smad1/5/8 (p-smad1/5/8) peaks at the mid-blastula stage in an immunoblot [81], following a similar trend as our proteomic dataset (Fig.5Gi). Smad 2 is a component of the Nodal pathway that mediates the specification of the oral ectoderm and suppresses formation of serotonergic neurons on the animal plate [96]. Smad 2 protein also peaks at the blastula stage and greatly diminishes into the gastrula and larval stages (Fig.5Gii). Smad4 protein is shown to increase dramatically from the blastula to gastrula stage, where it peaks in the larval stage (Fig.5Giii). Little is known about the function of Smad4 in sea urchin development; however, it is critical in other embryonic contexts, such as mouse cardiac development and neurogenesis [97] and zebrafish skeletal and cardiac muscle formation [98]. Due to its rise in relative protein levels in the gastrula and larval stages, Smad4 might play a conserved role in neurogenesis and muscle tissue formation in sea urchin development, similar to that of the mouse and zebrafish. Although proteomics data lack information on protein localization and protein phosphorylation states, the relative level of protein changes are consistent with the literature and can be informative.

Forkhead box (FOX) TFs are an evolutionarily conserved family of TFs that are critical in development and adult homeostasis [99]. Of the 22 Forkhead genes in the sea urchin genome [100], we identified FoxA, FoxO, Foxk1 and FoxJ3 (Fig.5H). FoxA is critical for gut formation [101], specifically maintenance of the endoderm–mesoderm boundary [102]. The expression of *FoxA* occurs during the blastula stage where it is lowly expressed in the veg2 endomesoderm [101, 103]. *FoxA* becomes asymmetrically restricted to the oral side of the endomesodermal ring [101] and eventually uniquely enriched in the endoderm of the gastrula. Our results indicate that FoxA protein increases in relative levels from 32-cell to blastula, similar with its mRNA expression pattern, and peaks at the larval stage (Fig.5Hi). Little is known about FoxO’s function in sea urchin development; however, it has been shown to regulate human embryonic stem cell pluripotency [104], T cell development and function [105], lens morphogenesis [106], and hematopoietic stem cell homeostasis [107]. The mRNA expression of *FoxO* is maternal and increases at the blastula stage, specifically in the skeletogenic cells and later in immune cells and the ciliary band [100]. The relative protein level of FoxO is similar to FoxA, in that it increases greatly from the 32-cell to blastula stage, and peaks at the larval stage (Fig.5Hi). It likely serves an immune or nervous system-related function due to its enriched mRNA expression in these cell types and the protein being highly present during later development (Fig.5Hi). The protein levels of FoxA and FoxO have a positive correlation with their corresponding mRNAs.

The two other Forkhead transcription factors identified (FoxK1 and FoxJ3) have high levels of maternal transcripts [100] (Fig.5Hii). The relative protein levels of both FoxK1 and FoxJ3 increase dramatically into the blastula stage, both peaking at the larval stage (Fig.5Hii). *FoxK1* transcript is highly expressed in the aboral ectoderm, skeletogenic cells, and later in immune cells of the gastrula, whereas *FoxJ3* is ubiquitously expressed throughout development [100]. Little is known of their specific functions in sea urchin development. The expression pattern of *FoxJ3* suggests that its maternal mRNA is degraded during the cleavage stage and undergoes zygotic transcription from the cleavage stage to the blastula stage. The FoxK1 protein accumulates steadily from the cleavage to the larval stage. From the blastula stage onward, the Foxk1 mRNA and protein levels have a positive correlation. In the case of FoxK1, its protein and mRNA levels are discordant; where its mRNA steadily decreases with its highest level in the egg and its protein increases steadily from the egg to the larval stage. Overall, the proteomics data capture some of the Fkh TF proteins throughout development.

### Proteins are positively correlated with their transcripts in differentiated immune and skeletogenic cells

These proteomics data can also be used to understand specific tissue development. The examples discussed here are the mesodermally-derived immune and skeletogenic cells. Mesodermally-derived cells give rise to a group of immune cells called pigment cells (PCs) and blastocoelar cells (BCs). Their differentiation is regulated by Delta/Notch and Nodal signaling pathways [77, 87, 108–110]. Delta/Notch signaling occurs in two waves, with the first wave important in activation of endomesodermal TFs in the posterior half of the embryo. Delta expression in the large micromeres (future PMCs) activates Notch in the neighboring cells (future PCs and BCs) to activate Gcm, a master regulator of non-skeletogenic mesoderm (NSM) [83, 108, 109, 111, 112]. We identified the relative Delta protein level to remain consistent from egg-32-cell, while its transcript peaks during the cleavage stage (Fig.6Ai). Relative Delta protein level thereafter decreases into the blastula and gastrula stages, while its transcript decreases into blastula but increases again in the gastrula and larval stages. Although average relative Delta protein increases from gastrula to larval stage, the deviation at larval stage is too great to gauge accurate level (Fig.6Ai). A caveat is that *Delta* is expressed in select few cells, and potentially for such proteins, proteomics data are limited in capturing their dynamics. After Nodal becomes restricted to the ventral (oral) ectoderm, it activates TF Not. Not signals neighboring vegetal tier 2 NSM cells at the ventral side of the embryo to become BCs via activation of BC-specific TFs Ese and Scl and inactivation of Gcm [83, 110]. Cells on the dorsal side of the NSM later become PCs. As stated previously, Nodal protein level increases into the 32-cell and blastula stage, likely reflecting its role in NSM differentiation (Fig.5Fi). We were unable to detect Not, Ese and Scl.

**Figure 6.**
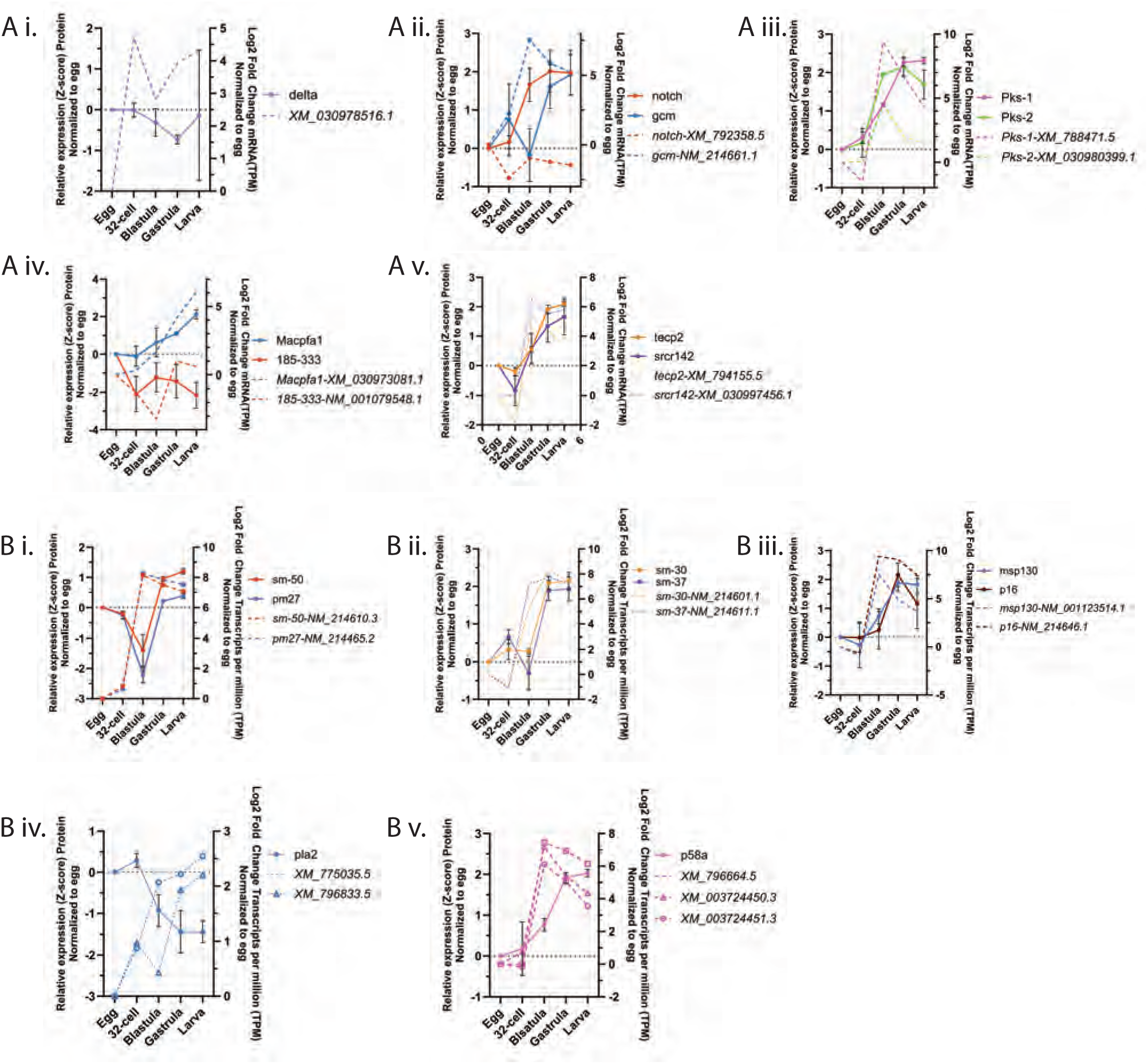
Proteomic analyses of Immune and skeletal cells. Proteins are denoted by solid lines and their corresponding mRNAs are of the same color denoted by dashed lines. (A) *Delta*, *Notch*, *Gcm*, *Pks-1*, *Pks-2*, *Macpfa1*, *185-333*, *tecp2* and *srcr142* mRNAs (Log_2_TPM) and proteins (Z-score) normalized to the egg are plotted. (B) *Sm-50*, *sm-27*, *sm-30*, *sm-37*, *msp-130*, *p16*, *pla2* and *p58a* mRNAs and proteins normalized to the egg are plotted.

The relative Notch protein expression increases dramatically from 32-cell to blastula stage, concurrent with its transcript increase and peaks at the gastrula and larval stage (Fig.6Aii). Although Delta/Notch signaling pathway transcriptionally activates *Gcm* [113], it does not appear that there is a definitive peak in Delta or Notch proteins preceding Gcm (Fig.6Ai, ii). However, Notch and Gcm both increase concurrently into the 32-cell stage, and Notch protein has the greatest fold-change increase prior to that of Gcm’s largest fold-change increase (Fig.6Aii). In contrast, Delta protein decreases from the cleavage stage to the gastrula stage and increases in the larval stage (Fig.6Ai). BCs, PCs, and multipotent progenitor cells are further differentiated during the second wave of Delta/Notch signaling during blastulation (17-22 hpf) [83, 108, 112, 114–116].

The PCs are fully specified when Gcm activates Pks1 around early blastula (14-16 hpf). PCs are the first of the NSM to ingress into the blastocoel at the mesenchyme blastula stage [109, 111, 117, 118]. PCs migrate to the ectoderm at the gastrula stage where they perform immune function [117, 118] or aid in wound repair together with BCs [119]. Pks1 (polyketide synthase 1) is a key enzyme expressed specifically in PCs, where they synthesize naphthoquinone pigments, such as echinochrome A with antimicrobial properties [120]. Pks2, which differs in its catalytic domains compared to Pks1, is expressed in PMCs, with unknown function [121]. Pks2 protein level has a 2- fold increase into blastula, while Pks1 protein level reaches 2-fold increase at the gastrula stage (Fig.6Aiii). The levels of transcript and protein of Delta and Notch that are more upstream in the immune cell specification and differentiation do not seem to correlate well. It appears that both the transcript and protein levels of differentiated BC and PC markers, such as Macpfa1, 185/333, tecp2, and srcr142 for BCs and Pks-1, Pks-2 for PCs, are more positively correlated (Fig.6Aiv, v). Thus, the protein and transcript levels of a signaling protein may be less correlative, while the transcript and protein levels of genes in terminally differentiated cell types are more positively correlated.

Sea urchin skeletogenesis is one of the most well studied processes of sea urchin development. The Delta/Notch pathway also plays a role in skeletogenic cell specification and differentiation [110]. The larval skeletal spicules are transient endoskeletal elements that are later abandoned during metamorphosis. PMCs form calcium carbonate spicules that function in protection, structural support, locomotion, and feeding orientation [112, 122–130]. PMCs are specified as early as the 16-cell stage as large micromere progenitors at the vegetal pole of the embryo [123, 131, 132].

Notable processes that the PMCs undergo are epithelial to mesenchymal transition from the vegetal plate and cell migration into the blastocoel of the embryo [133, 134]. Biomineralization occurs as PMCs fuse into a syncytium, where deposits of organic precursors of secreted organic matrix are made into the larval skeleton [127, 135]. Previous proteomic studies identified 231 proteins in the matrix of *S. purpuratus* skeletal spicules through their isolation from larvae using C_18_ reversed-phase liquid chromatography and mass spectrometry [24]. Using our novel in-cell proteomics E4 approach, we identified 143 of these 231 proteins (62%) using whole embryos without spicule isolation (Table S2). These included the spicule matrix and biomineralization proteins, including sm-30, sm-50, sm-37, msp-130, pm-27, p58a, p16 and pla2 (Fig.6Bi-v). In general, the transcript and protein levels of spicule matrix genes have similar trends with a positive correlation (except for pla2), where their mRNA levels peak at the blastula stage followed by protein level peaks at the gastrula and larval stages (Fig.6Bi-v). Because some mRNA and protein exhibit interesting positive and negative correlations (Figs.5, 6), we also examine the correlation between mRNA and protein levels across development for all genes identified in this study.

### Genome-wide correlation analysis between mRNA and protein levels reveals a majority of genes are negatively correlated

To examine the correlation between mRNA and protein levels during early embryogenesis, we cross-referenced all genes identified in our proteomics dataset (Table S1) with available transcriptomic data from the Echinobase database [18] (Table S2). After filtering our list of proteins for transcripts that have a TPM>0 in at least 1 timepoint from egg, 32-cell (10 hpf in transcript dataset), blastula, gastrula and larva stages, we obtained a list of 7,063 proteins (97% of total proteins) with corresponding mRNA counterparts (Table S2). We then performed a time resolved Pearson’s correlation analysis between the Log_2_ (iBAQ) from proteomics data and Log_2_ (TPM+1) transcriptomic data from egg-larva over developmental time from Echinobase (Fig.7A-C) [18, 136, 137]. We identified more proteins that have a strong negative correlation with their corresponding mRNA levels (r≤-0.5; 30% 2,117/7,063) than proteins that have a strong positive correlation with their corresponding mRNAs throughout development (r≥0.5) (23%1,620/7,063) (Fig.7A and Table S2). The global, mean correlation hovered near zero (r =-0.0645), indicating widespread temporal discordance between steady-state mRNA abundance and proteomic output (Fig.7A), similar to *Xenopus* early embryonic datasets and *Ciona* mRNA and protein correlation [136, 138]. For a majority of genes, protein levels are uncoupled from their corresponding mRNA expression levels in early sea urchin development. Positively correlated genes (n=1,620) are enriched in cytoskeletal organization, mitotic function and regulation of cell cycle (Fig.7Di), while negatively correlated genes (n=2,117) are enriched in gene expression and nucleic acid metabolism (Fig.7Dii). An example of a gene which has zero correlation is shown (Fig.7Diii). Global correlative analysis of mRNA and protein levels throughout development is lacking. However, a study across 14 human tissue types found a class of genes (1,012) that consistently show a highly positive correlation between mRNA and protein expression [139]. These genes encode proteins involve in oxidation/reduction and various transport activities, including ion transport and transmembrane transporter activity [139].

**Figure 7.**
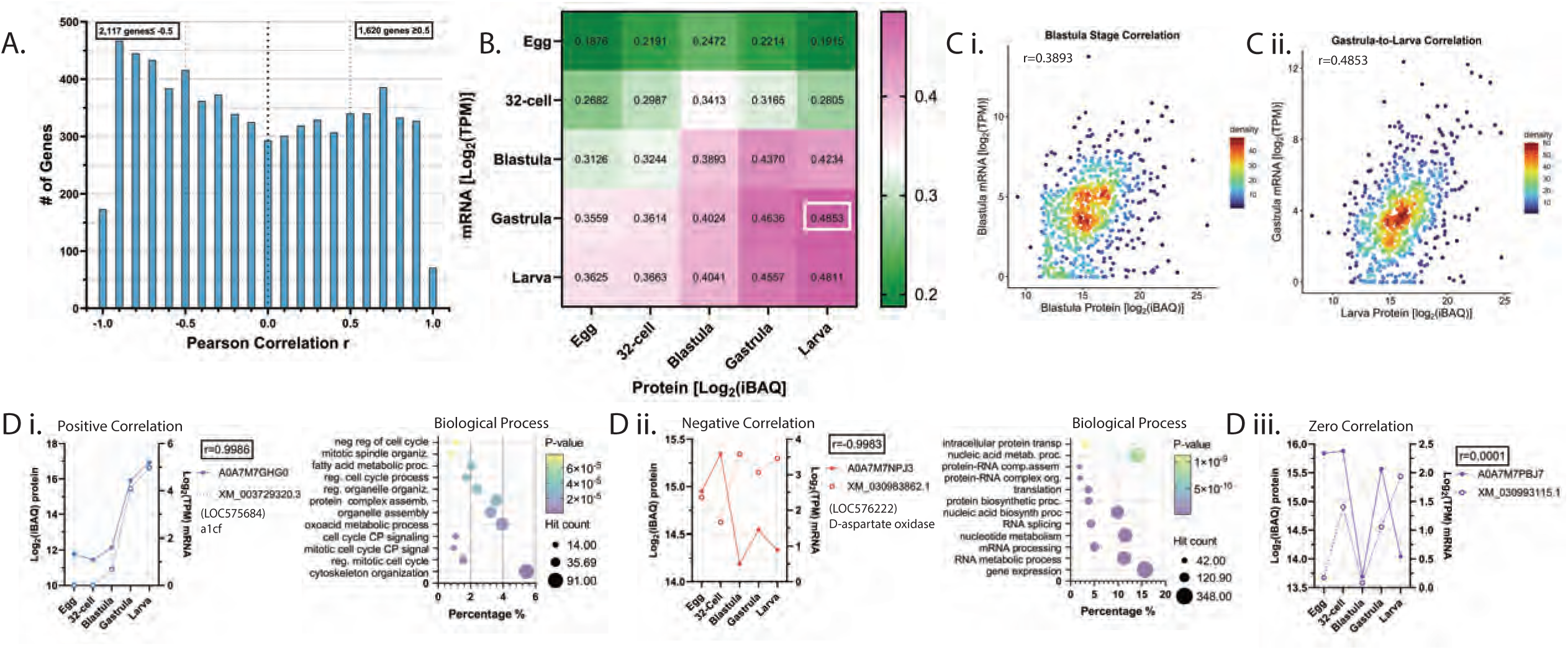
Genome-wide correlations between mRNA and protein levels. (A) Histogram of Pearson’s correlation distribution among mRNA and protein over development is plotted. 2,117 genes have r<-0.5 and 1,620 genes have r>0.5. (B) Heatmap of cross-gene correlation between mRNA and protein (7,063 genes) across developmental stages is depicted. Correlation coefficient (r) was calculated via Pearson’s correlation. Highest correlation is boxed. (C) Representative density plots of individual time points correlation between mRNA [Log2(TPM)] against protein [Log2(iBAQ)] are depicted. (Di) Example of highly positive correlated protein-mRNA. (Di) DAVID bioinformatics analysis (Biological Process) of significantly positive correlated genes. (Dii) Example of highly negative correlated protein-mRNA. (Dii) DAVID bioinformatics analysis (Biological Process) of significantly negative correlated genes. (Diii) An example of protein-mRNA with no correlation is depicted.

We then asked how well the abundance of the transcriptome predicts the abundance of the proteome at any given embryonic stage. A global cross-correlation matrix depicts the transition the embryo undergoes through development by comparing Log_2_ transformed mRNA (TPM) against protein (iBAQ) abundances across all five developmental stages (Fig.7B,C). Scatter plots (density-indicating the number of genes) of mRNA [Log_2_(TPM)] against protein [Log2(iBAQ)] abundances across all five developmental stages are shown (Fig.7C,S1C). A progressive linear tightening of transcript-protein coupling was observed along the diagonal, steadily increasing through the blastula (r=0.3893) and gastrula (r=0.4636) stages, and peaking at the larval stage (r=0.4811) (Fig.7B). Notably, the highest correlation in the entire matrix was captured between gastrula mRNA and larva protein (r=0.4853), revealing a prominent developmental time-lag wherein the increased transcripts in the gastrula stage are translated in the larval stage (Fig.7B,C). For example, comparing the mRNA-protein correlation in blastula vs. blastula stage, we observed correlation of r=0.3893 (Fig.7B,C). However, mRNA at blastula stage vs. proteins at gastrula stage has a stronger correlation of r=0.437 (Fig.7B,S1C). This observation of developmental time lag in mRNA and their protein level correlation is also exemplified in skeletal biomineralization protein-mRNA correlations (Fig.6Bi-v).

Overall, we present a novel in-cell proteomics approach that requires only 50 eggs/embryos to provide consistent and comprehensive proteomic information throughout sea urchin embryogenesis. The low input of embryos per sample makes this an attractive technique for gene perturbation studies that require microinjections or transplantations to execute. Further, our analysis revealed that most genes have discordant correlations between mRNAs and their corresponding protein levels. In addition, although mRNA quantity may peak at a certain developmental period, it may take longer for subsequent translation and accumulation of the protein later in development. In general, proteomes obtained from this study are consistent with literature, indicating utility of this data set as an important resource for the community.

## Figure Legends

**Figure S1.**
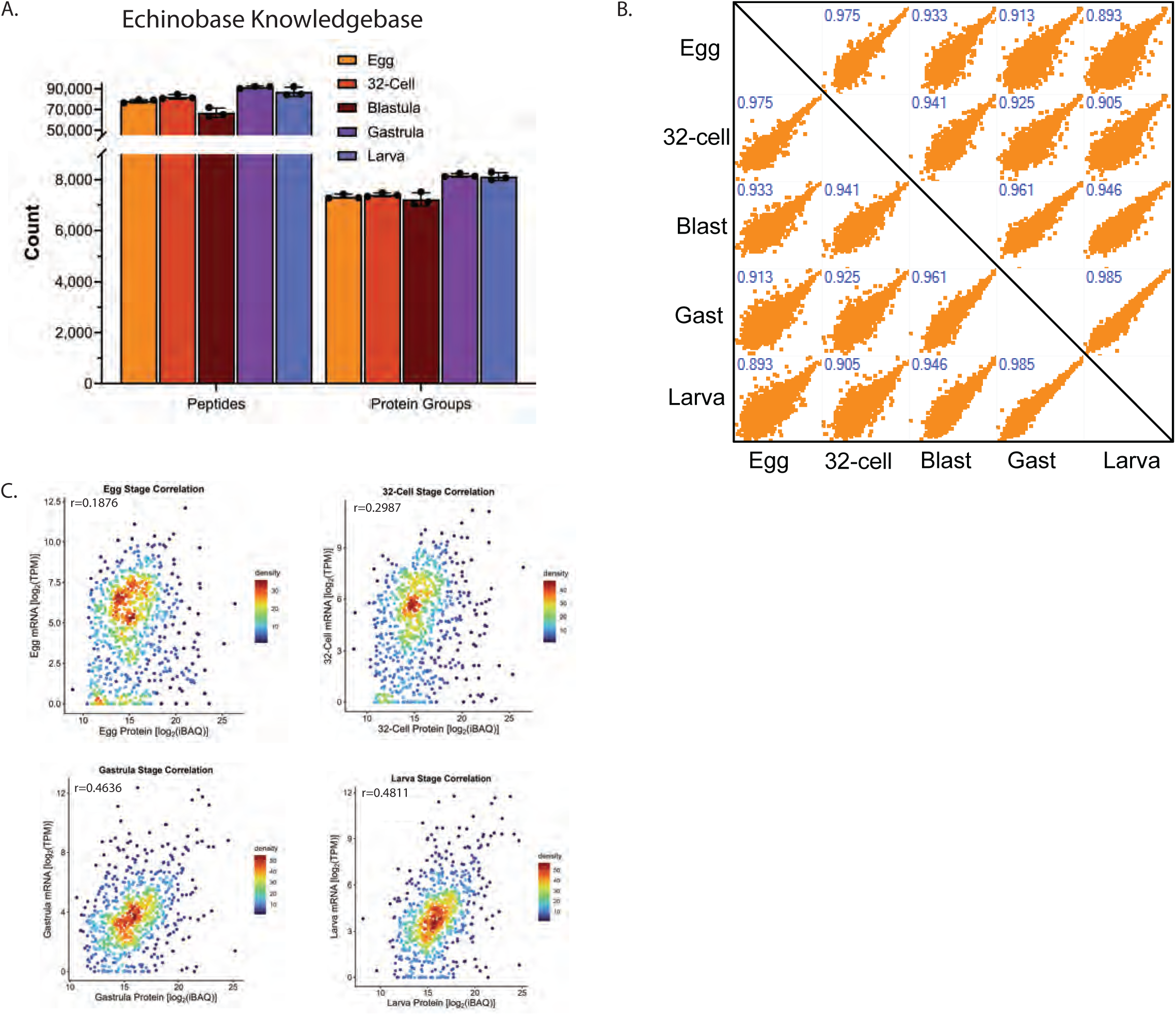
(A) Number of peptides and protein groups identified across 5 developmental stages (egg-larvae) using Echinobase is shown. Error bars represent standard deviation among 3 bioreps. (B) Spearman’s rank correlation across 5 developmental stages of log transformed protein abundances is shown. (C) Density plots of protein-mRNA correlations of all developmental time points are depicted.

